# Evidence for motility in 3.4 Gyr-old organic-walled microfossils ?

**DOI:** 10.1101/2020.05.19.103424

**Authors:** F. Delarue, S. Bernard, K. Sugitani, F. Robert, R. Tartèse, S.-V. Albers, R. Duhamel, S. Pont, S. Derenne

**Affiliations:** Sorbonne Université, CNRS, EPHE, PSL, UMR 7619 METIS, 4 place Jussieu, F-75005 Paris, France; Muséum National d’Histoire Naturelle, Sorbonne Université, UMR CNRS 7590, IRD, Institut de Minéralogie, de Physique des Matériaux et de Cosmochimie, IMPMC, 75005 Paris, France; Department of Earth and Environmental Sciences, Graduate School of Environmental Studies, Nagoya University, Nagoya, Japan; Department of Earth and Environmental Sciences, The University of Manchester, Manchester M13 9PL, United Kingdom; Molecular Biology of Archaea, Institute of Biology II, Faculty of Biology, University of Freiburg, Freiburg, Germany; Spemann Graduate School of Biology and Medicine, University of Freiburg, Freiburg, Germany

## Abstract

The oldest traces for planktonic lifestyle have been reported in ca. 3.4 billion years old silicified sediments from the Strelley Pool Formation in Western Australia. Observation of flange appendages suggests that Archean life motility was passive and driven by drifting of microorganisms in their surrounding environment. Until now, the oldest traces for active motility are ca. 2.1 billion years old. Whether or not active motility already existed during the Archean eon remains an open question. Here we report the discovery of new 3.4 billion years old tailed microfossils. These microfossils exhibit a lash-like appendage that likely provided them with movement capabilities. This suggests that these microfossils are the oldest remains of active motile life forms. With the ability to move in liquids and on organic and/or mineral surfaces, these microorganisms were capable of escaping from harsh environments and/or colonizing new ecological niches as early as 3.4 billion years ago. The existence of these deep-rooted Archean motile life forms offers a new picture of the Archean biodiversity, with unanticipated evolutionary innovative morphological complexities.

## Introduction

Archean carbonaceous microfossils illustrate the widespread presence of life on Earth as early as ca. 3.4 billion years ago (Westall et al., 2006; Sugitani et al., 2010; Wacey et al., 2011; Alleon et al., 2018; Delarue et al., 2020). However, the interpretation of the Archean palaeobiological record is fraught with difficulties pertaining to fossilization and burial-induced degradation processes, as illustrated by intense debates over the past couple of decades (Schopf et al., 2002; Brasier et al., 2002; Wacey et al., 2016)). Remnants of early life forms all have experienced burial and thermal alteration for billions of years, which led to the degradation of many pristine biological traits (Javaux et al., 2019). Therefore, Archean putative microfossils tend to exhibit simple morphological shapes (e.g., spheroidal, filamentous, film, and lenticular forms) that can also be abiotically produced (Garcia-Ruiz et al., 2003; Cosmidis et al., 2016), precluding, in turn, any simple morphological distinction between genuine biological remnants and mineral/organic biomorphs. Because of the lack of taxonomically informative features (Javaux et al., 2019), morphological criteria alone are generally considered as insufficient to assess the biological nature of ancient traces of life in the Archean geological record (Brasier et al., 2006). As a result, the ancient fossil record has not yet conveyed a complete picture of ancient biodiversity. Here, we report the discovery of 3.4 billion years old organic microfossils from the Strelley Pool Formation (SPF) from Western Australia exhibiting exceptionally preserved morphological traits indicative of active motility.

## Results and Discussion

Observations of thin sections of SPF reveal the existence of tailed organic-walled microfossils (Fig. 1a, b). These tailed organic-walled microfossils are exclusively observed within the main siliceous sedimentary matrix precluding their introduction during hydrothermal fluid circulation post 3.4 Ga. Raman spectra of tailed specimens chemically isolated from the mineral matrix are typical of those of disordered carbonaceous materials having undergone a low-grade metamorphism (Fig. 2a; Pasteris and Wopenka, 2003; Delarue et al., 2016). Their Raman line shapes suggest that these microfossils experienced peak temperatures of approximately 250-300 °C (Lahfid et al., 2010). Raman first-order spectra of studied SPF tailed microfossils are similar to those previously determined on syngenetic microfossils from the same geological formation observed in thin sections (Lepot et al., 2013; Sugitani et al., 2013), on freshly fractured faces (Alleon et al., 2018), and in acid maceration residue (Delarue et al., 2020). Therefore, these tailed organic-walled microfossils should be regarded as syngenetic as they were subjected to the maximum metamorphic temperature registered by their host rock.

**Figure 1:**
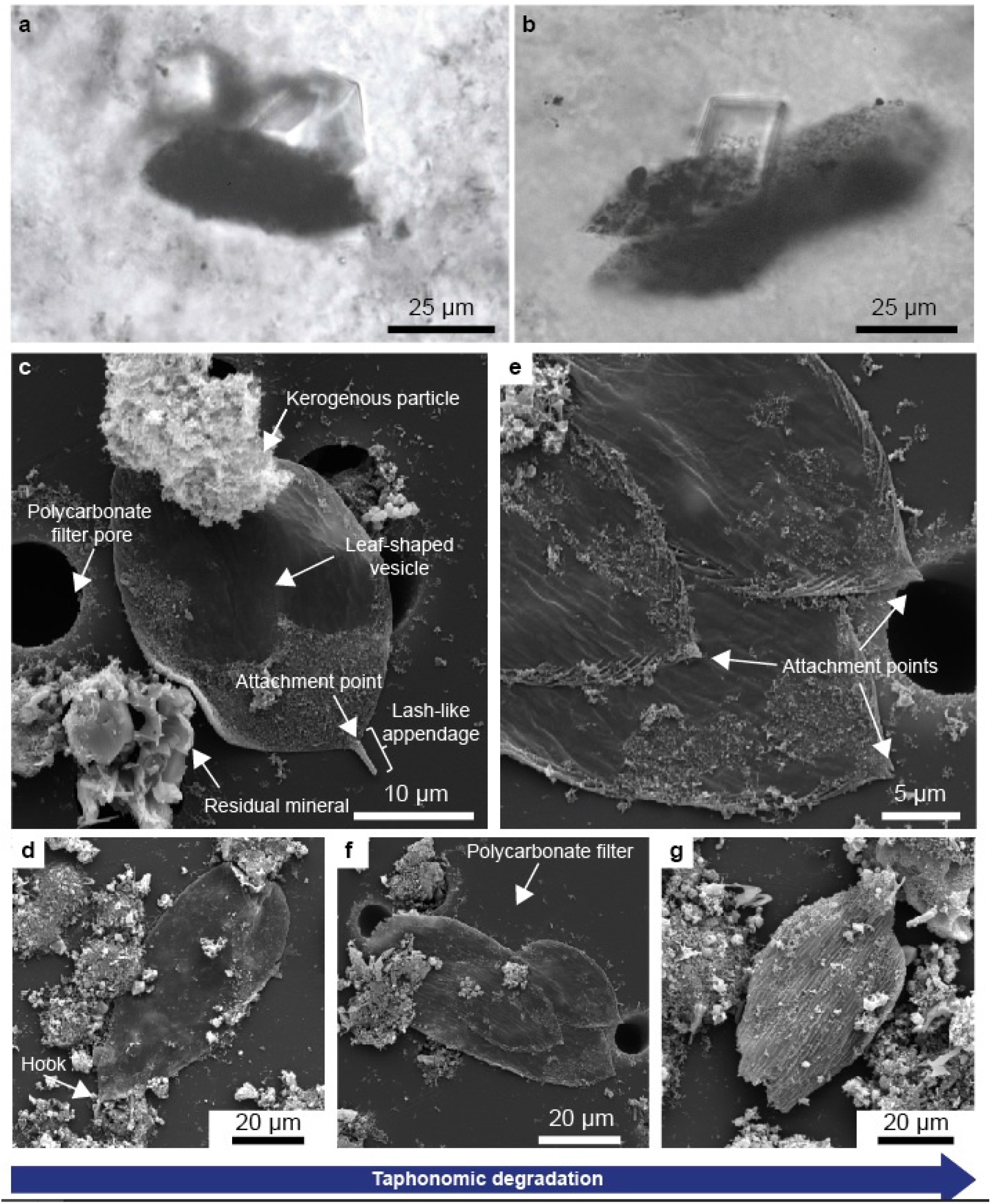
Thin section micrographs and scanning electron microscopy images of tailed organic microfossils. (a-b) Micrographs presenting two specimens embedded in the main mineral matrix of the studied SPF chert. (c-f) SEM images of tailed organic microfossils isolated by acid maceration. (c,d) Exceptionally-well preserved leaf-shaped vesicles presenting a locomotory organelle composed of an attachment point and of a lash-like appendage (e-g) Corresponding degraded organic-walled microfossils. A taphonomic degradation gradient is observed from the left to the right. Classic taphonomical degradation features, including folds and tears, are observed.

**Figure 2:**
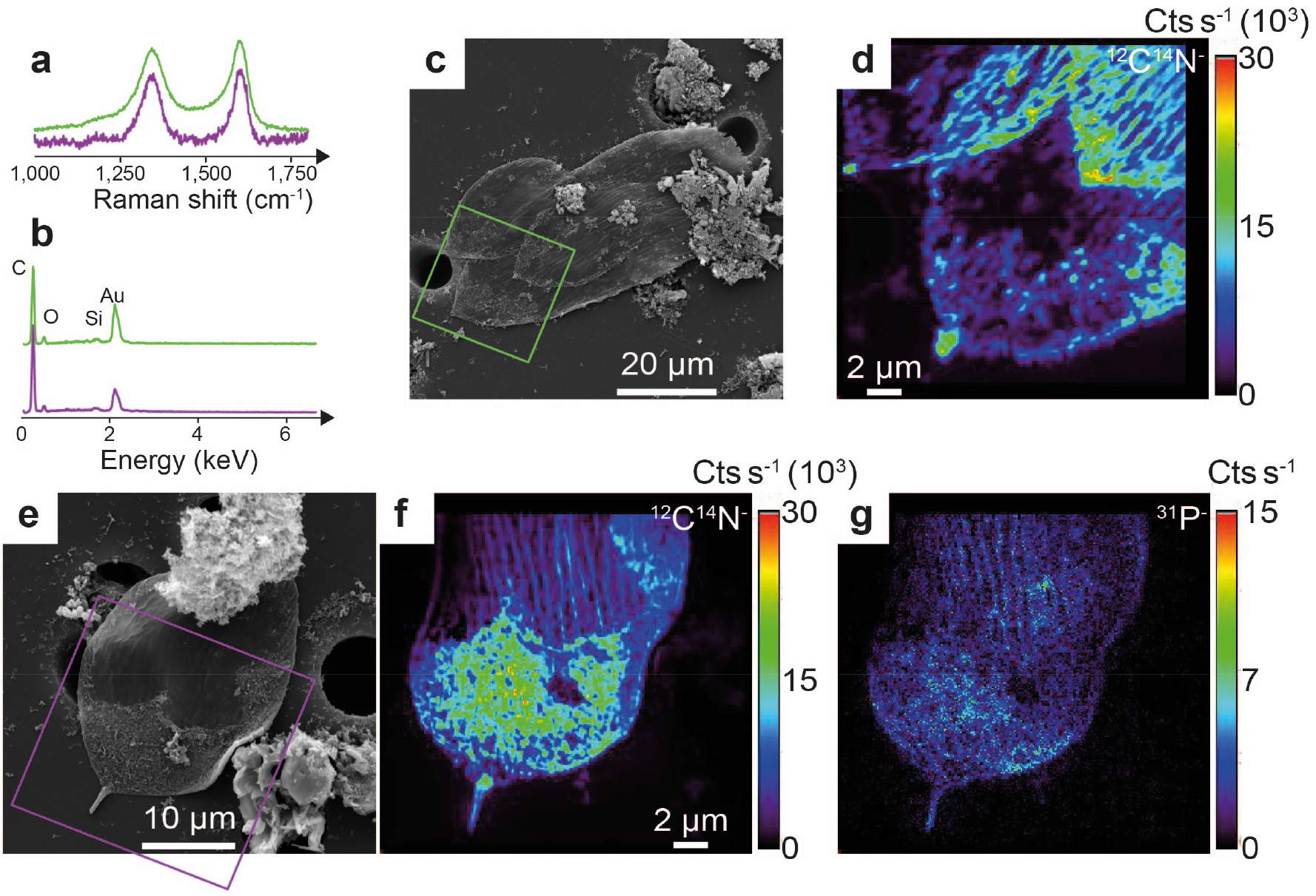
Raman spectra, energy-dispersive X-ray spectra and nanoscale secondary ion mass spectrometry images. (a) First-order Raman spectra determined on isolated tailed-organic walled microfossils and (b) corresponding energy-dispersive X-ray spectra. Green and purple lines indicate that spectra were acquired on specimens shown in panels c and e, respectively. (c, e) SEM images of organic-walled microfossils investigated by EDX, Raman spectroscopy and NanoSIMS. Green and purple squares indicate areas probed by NanoSIMS. (d, f) the ^12^C^14^N^-^ ion images illustrate the presence of nitrogen. (g) the ^31^P^-^ image illustrates significant levels of phosphorus. No significant level of ^31^P^-^ was recorded on the second specimen shown in panel c.

If Raman spectroscopy is a useful tool to assess syngeneity, it is not sufficient to determine the biogenicity of putative remains of ancient life (Pasteris and Wopenka, 2003). Energy-dispersive X-ray spectroscopy data show that the studied specimens essentially contain C and O (Fig. 2b), confirming their organic nature, while nanoscale secondary ion mass spectrometry reveals significant levels of nitrogen and, in one specimen, phosphorus (Figs. 2d, f-g). The presence of these key elements of cell walls, proteins, and nucleic acids are consistent with a biological origin. Spatially resolved chemical investigations exploiting X-ray absorption confirm the heterogeneous chemical nature of the investigated organic-walled microfossils as at least three different chemical structures could be distinguished in a given specimen (Fig. 3). Specimens contain some highly graphitic organic materials with almost no nitrogen and a X-ray absorption spectrum exhibiting a broad peak of conjugated aromatic groups (285.5 eV) and the excitonic absorption feature of planar domains of highly conjugated π systems (291.7 eV; Bernard et al., 2010). Closely associated are N-poor materials with XANES spectra similar to those of thermally-altered kerogen with an intense absorption peak at 285 eV (aromatic or olefinic groups), a relatively broad absorption feature at 287.5 eV (aliphatic carbons), and an absorption feature at 286.6 eV (imine, nitrile, carbonyl and/or phenol groups; Bernard et al., 2010; Le Guillou et al., 2018). Specimens also contain N-rich compounds (N/C ~ 0.22) with XANES spectra that exhibit clear contributions of quinones or cyclic amides (284.5 eV), aromatic or olefinic carbons (285.1 eV), imine, nitrile, carbonyl and/or phenol groups (286.6 eV), aliphatics (287.7 eV) and amides (288.2 eV). Altogether, the chemical structure of the SPF specimen investigated here is consistent with the preservation of partially degraded biomolecules.

**Figure 3:**
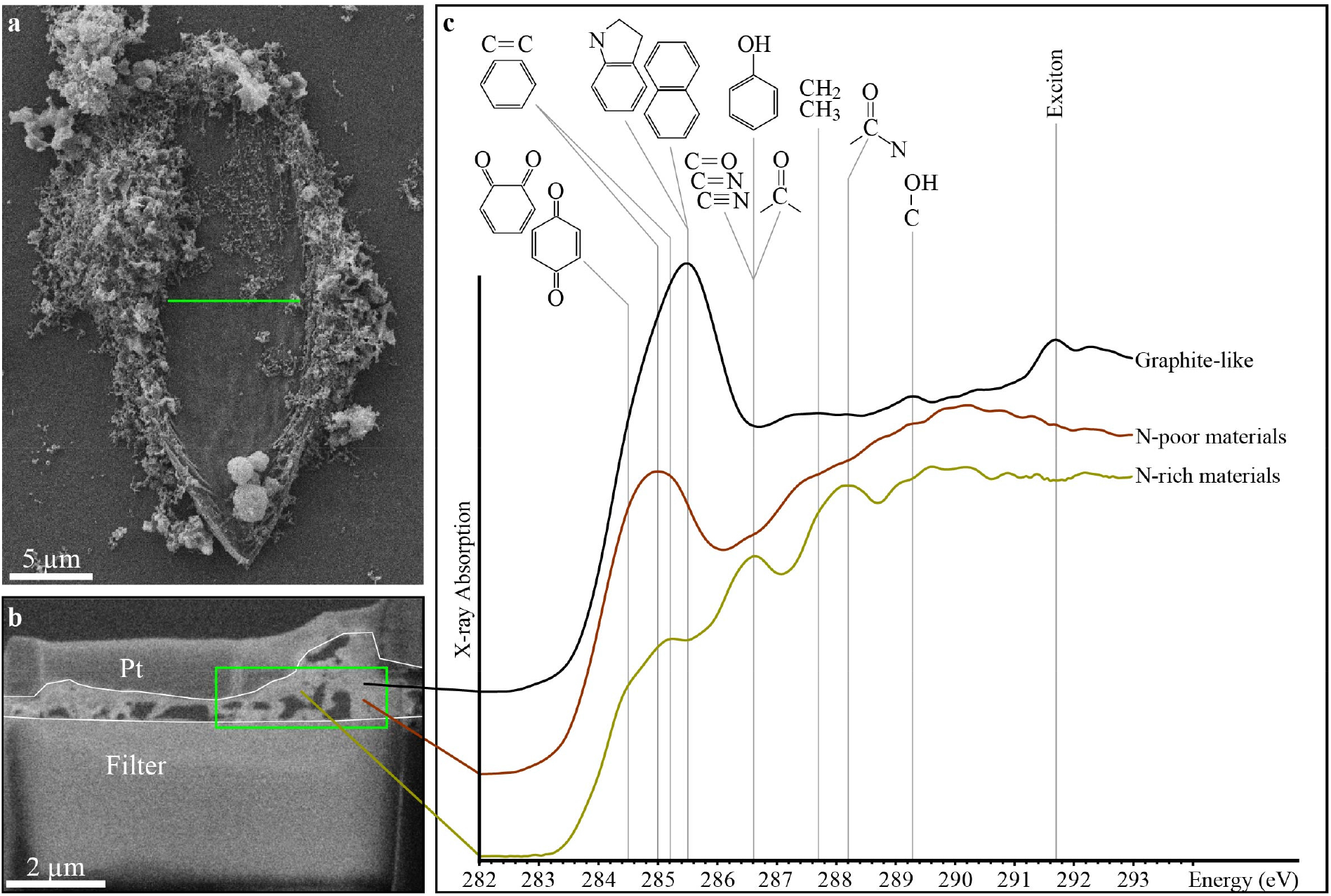
Scanning transmission X-ray microscopy -based X-ray absorption near edge structure characterization. (a) SEM image of the specimen from which a focused ion beam foil has been extracted (green line). (b) SEM image of the focused ion beam foil evidencing the low thickness of the specimen. The green square indicates the area investigated using STXM. c, Carbon - X-ray absorption near edge structure spectra of the organic materials composing the investigated specimen.

From a morphological point of view, the organic-walled microfossils are leaf-shaped vesicles ranging from 30 to 84 μm in length and from 16 to 37 μm in width (Fig. 1). They exhibit classic taphonomical degradation features, including folds and tears (Figs. 1c-g). The preparation of ultrathin foils using focused ion beam illustrates their relative limited thickness, ranging from 200 to 500 nm (Fig. 3). Some specimens also exhibit a specific morphological feature, a lash-like appendage protruding from the leaf-shaped cell (Figs. 1c, d).

Based on analogous morphological traits observed in past and in modern microorganisms, several origins/functions may be hypothesised to explain the occurrence of this lash-like appendage. A number of extant microorganisms exhibit a micrometric tube-like appendage called prostheca, which can be involved in anchoring cells to organic and mineral surfaces, in nutrient uptake or in asexual reproduction by budding at its tip (Curtis, 2017). To assess whether the observed lash-like appendage is a remnant of an ancient prosthecum, we propose an Appendage Shape Index (ASI) based on the ratio between the width of the appendage and of the parent cell (Fig. 4). Compilation of morphometric data on extant microorganisms suggests that ASI ranges between 15 and 45% in prostheca, while the lash-like appendages observed in SPF microfossils are characterized by ASI ranging between 2.2 and 5.8 %, consistently with those observed on modern archaella, flagella and cilia (Fig. 4). In addition, a prosthecum consists in an extension of the cellular membrane, implying a structural continuity between the microorganism body and the base of the prosthecum (Javaux *et al*., 2003). On the studied SPF specimens, we observed an anchoring attachment point and a filament-like appendage, indicative of two distinct structural subunits (Fig. 1). Based on these morphometric and structural features, the lash-like appendages observed in SPF microfossils cannot be considered as remnants of a prosthecum. As far as we are aware, and in accordance with their ASI (Fig. 4), such distinct external and functional subunits can only be assigned to locomotory organelles. However, the lash-like appendages observed in SPF microfossils are between 0.7 and 1.2 μm in diameter, which is much larger than those reported for archaella, flagella, and cilia reaching ca. 10, 20, and 200 nm, respectively (Jarell and McBride, 2008; Beeby et al., 2020). Large cell dimensions (Ø > 10 μm) is a morphological feature commonly observed in Precambrian organic-walled microfossils (Javaux et al., 2010; Sugitani et al., 2010; 2015; Balidukay et al., 2016; Loron et al., 2019). Overall, the consistency of ASI values for SPF microfossils compared to those for archaella, flagella and cilia suggests that proportions between cell size and functional morphological traits/organelles may have persisted over the Earth’s history. However, the SPF microfossils’ lash-like appendages do not meet standard structural features (for instance, a curved hook connecting the filament to the basal body in flagella) observed on locomotory organelles from any organism of the three extant domains of life (see Khan and Scholey, 2018). Therefore, we propose that the lash-like appendage observed in some SPF microfossils may be a remain of a proto-locomotory organelle from a common ancestor or, alternatively, of an unknown and extinct domain of life.

**Figure 4:**
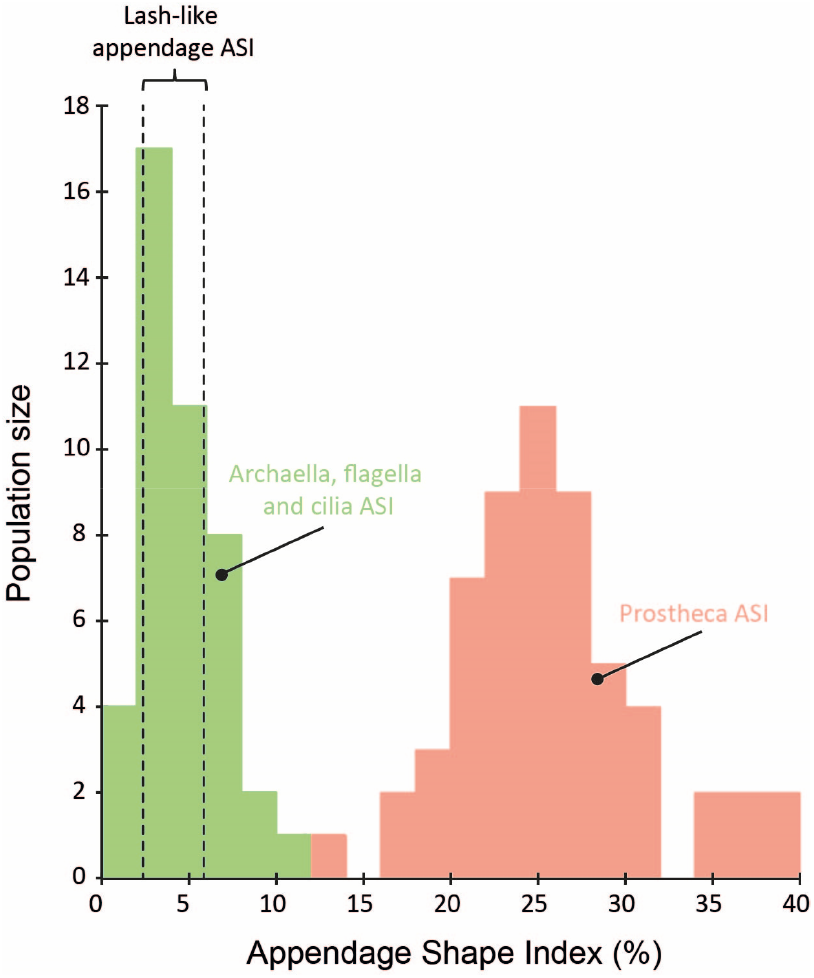
Compilation of Appendage Shape Indices determined on extant microorganisms. ASI was computed according to the ratio between the width of appendage (archaellum, flagellum, cilium and prosthecum) and of its parent cell (×100). Each width of appendage and of its parent cell was determined graphically following micrographs and images previously published in Southam et al. (1990), Poindexter and Staley (1996), Furuno et al. (1997), Wustman et al. (1997), Qintero et al. (1998), Wang et al. (2001), Miller et al. (2004), Bergholtz et al. (2006), Vasilyeva et al. (2006), Wagner et al. (2006), Kanbe et al. (2007), Abraham et al. (2008), Nge et al. (2008), Pyatibratov et al. (2008), Siano et al. (2009), Craveiro et al. (2010), Wang et al. (2011), Abraham and Rohde (2014), Chang Lim et al. (2014), Albers and Jarrell (2015), Deng et al. (2016), Kinosita and Nishizaka (2016), Sugitomo et al. (2016), Curtis (2017), Leander et al. (2017). ASI determined on archaella, flagella and cilia are indicated in green while those determined on prostheca are indicated in pink. The area delimited by dotted lines indicate ASI determined on four lash-like appendages observed on tailed SPF organic-walled microfossils. ASI ranges from 4.8 to 5.8 % and from 2.2 to 3.3 % in organic-walled microfossil observed in thin sections (n = 2) and in the acid maceration residue (n = 2), respectively. ASI is likely overestimated in thin sections as a consequence of shadows occurring at the edge of organic-walled microfossils.

Previous observations of 3.4-3.0 billion years-old flanged microfossils implied passive motility, microbial planktons drifting depending on its surrounding environment to engender movement (House et al., 2013; Sugitani et al., 2015; Oehler et al., 2017; Kozawa et al., 2019). To date, the oldest evidence for active motility was recorded as tubular sedimentary structures in 2.1 billion years old Francevillian sedimentary series (Gabon, El Albani et al., 2019). The preservation of lash-like appendages in some SPF microfossils suggests that some microorganisms were capable of active motility - a mechanism whereby microorganisms can direct where they go (Pollitt and Diggle, 2017) - as early as 3.4 Gyr ago. Since it likely provided them with the ability to move in the water column or at the surface of organic and/or mineral surfaces, this evolutionary morphological innovation marks a major step in the history of life on Earth, and provide a more complex picture of the Archean biodiversity. It suggests that microorganisms were already able to escape harsh environments, adapt their feeding strategies moving towards more favorable nutrient sources, and colonize new ecological niches less than a billion years after the Earth became habitable (Javaux et al., 2019).

## Methods

### Chemical isolation of microfossils

Organic-walled microfossils were isolated from the SPF carbonaceous black chert sample using a modified version of the classical acid maceration procedure (Delarue et al., 2020). A ‘soft’ acid maceration procedure was applied in order to minimize both potential physical and chemical degradations of organic microstructures. Prior to acid maceration, about 30 g of rock samples were fragmented into ~3 g rock chips rather than crushed into finer grains. Rock chips were cleaned using ultrapure water and a mixture of dichloromethane/methanol (v/v: 2/1). Rock chips were then directly placed in a Teflon vessel filled with a mixture of HF (40%, reagent grade) /HCl (37%; reagent grade; v/v: 9/1) at room temperature. After 48 hours, successive centrifugation and rinsing steps using ultrapure water were performed until reaching neutrality. The residual material was suspended in ethanol and filtered on polycarbonate filters (pore Ø = 10 μm). After ethanol evaporation, polycarbonate filters were fixed on carbon tape and coated with 20 nm of gold to prevent further contamination by atmospheric deposits and further analyses with SEM-EDXS and NanoSIMS

### Scanning electron microscopy and Energy Dispersive X-Ray Spectroscopy (SEM-EDXS)

SEM-EDXS imaging and analysis were performed on gold-coated filters using a TESCAN VEGA II at the French National Museum of Natural History (MNHN) operated with an accelerating voltage of 15 kV.

### Raman spectroscopy

Raman microspectroscopy was carried out using a Renishaw InVIA microspectrometer equipped with a 532 nm green laser. The laser was focused on the sample by using a DMLM Leica microscope with a 50× objective. The spectrometer was first calibrated with a silicon standard before the analytical session. For each target, we determined the Raman shift intensity in the 1000 to 2000 cm^-1^ spectral window that includes the first-order defect (D) and graphite (G) peaks. A laser power below 1 mW was used to prevent any thermal alteration during spectrum acquisition. Spectra acquisition was achieved after three iterations using a time exposure of 10 seconds. Raman microspectroscopy was performed on gold-coated organic surfaces implying a slight lowering of the D bands in comparison to the G one (see Delarue et al. 2020 for details)

### Nanoscale secondary ion mass spectrometry

Isolated microfossils were analyzed using a CAMECA NanoSIMS 50 ion probe using a Cs+ primary ion beam. Before measurements, pre-sputtering was performed over 30 × 30 μm2 areas for ca. 8 minutes using a 500 pA primary current (750 μm aperture diaphragm) to avoid surficial contamination, and achieve Cs+ saturation fluence and constant secondary ion count rates. Analyses were then carried out using a 10 pA primary current (200 μm aperture diaphragm) on smaller areas to avoid pre-sputtering edge artifacts. The secondary molecular species ^12^C^14^N^-^ and ^31^P^-^ were collected simultaneously in electron multipliers. The NanoSIMS raw data were corrected for a 44 ns dead time on each electron multiplier and processed using the Limage software.

### Focused ion beam (FIB)

FIB ultrathin sections were extracted from the organic microfossils using an FEI Strata DB 235 (IEMN, Lille, France). Milling at low gallium ion currents minimizes common artefacts including: local gallium implantation, mixing of components, creation of vacancies or interstitials, creation of amorphous layers, redeposition of the sputtered material on the sample surface and significant changes in the speciation of carbon-based polymers.

### Scanning transmission X-ray microscopy

XANES investigations were conducted using the HERMES STXM beamline at the synchrotron SOLEIL (Gif-sur-Yvette, France). Carbon contamination on beamline optics was constantly removed thanks to a continuous flow of pure O2. The well-resolved 3p Rydberg peak of gaseous CO2 at 294.96 eV was used for energy calibration. Collecting image stacks at energy increments of 0.1 eV with a dwell time of ≤ 1 ms per pixel prevented irradiation damage. The estimations of N/C values and the normalization of the C-XANES spectra shown here were done using QUANTORXS (Bernard et al., 2010).

## Acknowledgments

We thank V. Rouchon and O. Belhadj (CRCC) for Raman spectroscopy and D. Troadec (IEMN) for FIB extraction.. We also acknowledge The National NanoSIMS Facility at the MNHN, supported by MNHN, CNRS, Région Ile de France, and Ministère de l’Enseignement Supérieur et de la Recherche. Special thanks go to Stefan Stanescu and Sufal Swaraj for their expert support with the HERMES STXM beamline at SOLEIL. The HERMES beamline (SOLEIL) is supported by the CNRS, the CEA, the Region Ile de France, the Departmental Council of Essonne and the Region Centre. This work was supported by the Programme National de Planétologie (PNP) of CNRS/INSU, co-funded by CNES R.T. also acknowledges the UK Science and Technology Facilities Council (grant ST/P005225/1) for financial support.

## Authors contributions

F.D. designed the study and supervised most of the analyses. K.S. provided the samples investigated. S.B. conducted STXM-XANES analyses. R.D. conducted NanoSIMS analyses. S.P. conducted SEM-EDXS analyses. F.D. wrote the present manuscript with critical inputs from all authors.

